# Not just methods: User expertise explains the variability of outcomes of genome-wide studies

**DOI:** 10.1101/055046

**Authors:** Katie E. Lotterhos, Olivier François, Michael G.B. Blum

**Author notes:** Corresponding author Michael Blum Laboratoire TIMC-IMAG, Faculté de Médecine, 38706 La Tronche, France.

## Abstract

Genome scan approaches promise to map genomic regions involved in adaptation of individuals to their environment. Outcomes of genome scans have been shown to depend on several factors including the underlying demography, the adaptive scenario, and the software or method used. We took advantage of a pedagogical experiment carried out during a summer school to explore the effect of an unexplored source of variability, which is the degree of user expertise.Participants were asked to analyze three simulated data challenges with methods presented during the summer school. In addition to submitting lists, participants evaluated a priori their level of expertise. We measured the quality of each genome scan analysis by computing a score that depends on false discovery rate and statistical power. In an easy and a difficult challenge, less advanced participants obtained similar scores compared to advanced ones, demonstrating that participants with little background in genome scan methods were able to learn how to use complex software after short introductory tutorials. However, in a challenge of intermediate difficulty, advanced participants obtained better scores. To explain the difference, we introduce a probabilistic model that shows that a larger variation in scores is expected for SNPs of intermediate difficulty of detection. We conclude that practitioners should develop their statistical and computational expertise to follow the development of complex methods. To encourage training, we release the website of the summer school where users can submit lists of candidate loci, which will be scored and compared to the scores obtained by previous users.

## Introduction

The possibility to obtain dense genomic markers at reasonable cost has served the ambition to map genomic regions involved in biological adaptation (Schoville *et al.* 2012, Savolainen *et al.* 2013, Haasl and Payseur 2016). Various statistical methods and software have been developed to fulfill this ambition (Rellstab *et al.* 2015, FranÇois et al. 2016). The performances of these methods have been evaluated and compared under various evolutionary scenarios (Narum and Hess 2011, De Mita et al. 2013, De Villemereuil et al. 2014, Lotterhos and Whitlock 2014, Lotterhos and Whitlock 2015). Results obtained with genome scan approaches have been found to depend on the demographic history, on the genetic architecture of adaptation, on sampling design, on statistical software, with interactions between these factors. However, an unexplored source of variability in the application of genome-scan methods is the extent to which different users can achieve different outcomes. In this study, we report the results of a pedagogical experiment carried out at the “Software and Statistical Methods in Population Genomics” 2015 summer school (SSMPG 2015) to measure how the outcomes of genome scan methods vary among users having different prior levels of expertise.

## Material and Methods

The objective of the summer school held in September 2015 was to teach a set of recently developed genome scan methods for the detection of genomic regions involved in local adaptation. The teaching process was based on active learning in which participants were asked to perform data analyses of simulated data using the methods presented (Freeman et al. 2014). Three distinct challenges were proposed to the participants who had no *a priori* knowledge of the loci simulated under selection in each challenge. For each challenge, the participants could download simulated genomic data (SNPs) from a dedicated website. They were asked to submit lists of candidate SNPs using the methods presented during the teaching sessions. Lists built from combinations of methods were accepted.

The data for the challenges were simulated before the summer school by one instructor, who was the only person who knew how the data were simulated.A total of 48 attendees and the 5other instructors participated in analyzing the challenges. The datasets contained simulated genotypes for a fictive species. A vast majority of the simulated loci corresponded to selectively neutral alleles while a small fraction of them corresponded to adaptive alleles. The data were simulated by using the computer program NEMO (Guillaume and Rougement 2006).

The demographic history of the fictive species corresponded to a two-refugia model in a mountain range with three peaks. The species was initially limited to two nunatak (mountain top refuges) during a glaciation period of 3000 generations (Supp Fig 1). At generation 3000,a third nunatak was colonized because of climate warming. At generation 4000, all populations were colonized. For all three challenges, populations had the same demographic history of carrying capacity. However, migration rates or genetic architecture were different between challenges to create increasing difficulty for detecting selection (Supp Fig 2). For all simulations, neutral loci and Quantitative Trait Loci (QTL) were simulated on a genetic map of 6 linkage groups, each 100 cM long, with a resolution of 1 cM. Simulations of the first challenge assume 12 unlinked QTLs of equal effects on the trait and an island model of migration. Simulations of the second challenge had the same 12 QTLs of equal effects but migration declined with distance and there was less time until sampling. Simulations of the second and third challenge used the same values of migration rates. In the third challenge, there were 36 QTLs with effects on the trait as well as some linkage among QTLs. The parameter files used to create the simulations are included as supplementary material (SI Files 1-3).

Six statistical methods were presented to the participants during 45-minute teaching sessions. The statistical methods were divided into two categories: population differentiation (PD) methods and ecological association (EA) methods. PD methods use allele frequencies to compare single-locus estimates of a population differentiation statistic with their expectation from a null model. EA methods seek for genetic markers significantly correlated with one or several ecological or environmental variables (Rellstab et al. 2015). Methods based on PD methods included HapFlk (Bonhomme et al. 2010, Fariello et al. 2013), OutFLANK (Whitlock and Lotterhos 2015), pcadapt (Duforet-Frebourg et al. 2016), and SelEstim (Vitalis et al. 2014).EA methods included BayPass (Gautier 2015) and LFMM as implemented in the R package LEA (Frichot et al. 2013, Frichot and François 2015).

For each challenge, participants submitted lists of candidate loci and each list was evaluated by a score. We used an F-score that accounts for sensitivity (or power, POW) and falsediscovery rate (FDR) as follows (Fawcett 2004):

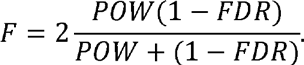

FDR was defined as the proportion of neutral loci in the list and power was defined as the proportion of adaptive loci contained in the list. *F*-scores range between 0 and 1. The *F*-score was equal to 0 for lists that contained no adaptive loci, and it was equal to 1 if and only if a list matched the list of truly adaptive loci perfectly. Using *F*-scores, we were able to evaluate the performances of the participants when using a particular method, as well as the variability of their performances in each challenge. At SSMPG 2015, most participants evaluated the number of loci under selection in the simulated datasets quite accurately, although their candidate lists may not have contained true positives only. Under this condition, the *F*-score was mainly an evaluation of the power (POW) of the participant's approach to detect truly adaptive loci. In this study, our intention is to understand the variability in F-scores of candidate lists submitted by users, rather than variability in the *F*-scores among the programs employed to detect adaptation.

For all challenges, participants were able to share expertise and collaborate with each other by building teams. When submitting lists of candidate loci to the website, users (a team or a single individual) had to declare their level of expertise. Two levels of expertise were predefined as “advanced user” or “less-advanced user”. For each challenge, a different number of users submitted lists and declared themselves as “advanced users” or “less-advanced users”, respectively. In the first challenge, there were 7 advanced and 67 less-advanced submissions, in the second challenge, there were 10 advanced and 56 less-advanced submissions, and in the third challenge, there were 10 advanced and 36 less-advanced submissions.

## Results

### Difficulty levels of challenges

To provide evidence that the difficulty of correctly identifying adaptive loci increased for each challenge, we evaluated the distribution of scores for each challenge (Figure 1).The median of the scores decreased with the challenge number. The median score (± standard deviation) was equal to 0.96 (± 0.16) forthe first challenge, 0.40 (± 0.24) for the second challenge, and 0.31 (± 0.24)for the third challenge. The first case was an easy challenge proposed to all attendees to test software installation and the online submission process. Participants focused their efforts on the challenges 2 and 3. In the following paragraphs, we report results obtained for the second and third challenges.

**Fig 1:**
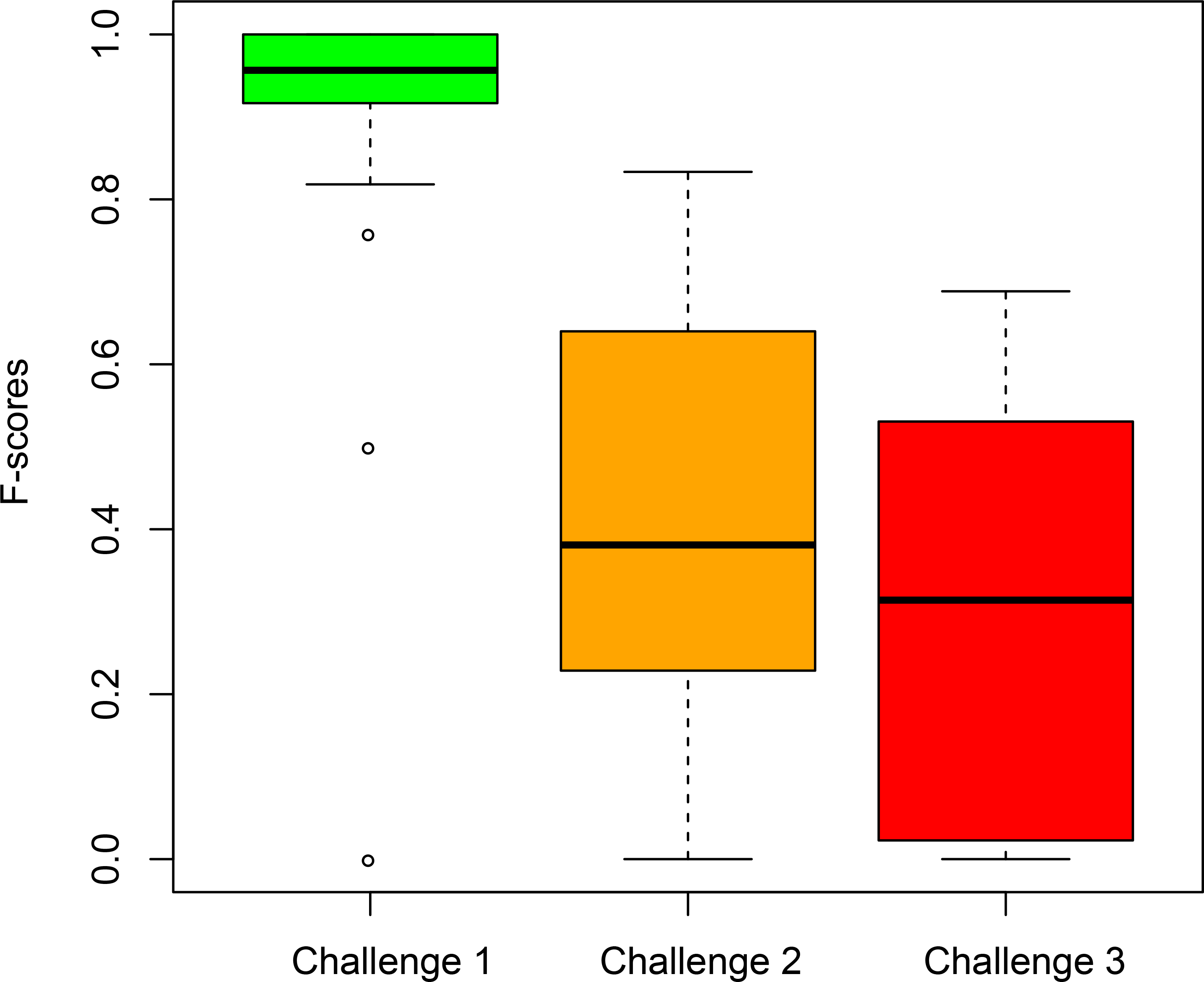
Distribution of F-scores for the three different data challenges.

## Software usage

For the second and third challenges, 124 candidate lists were submitted to the website and 111 submissions were retained after filtering for obvious errors or handling mistakes. The distribution of software usage was almost balanced (Figure 2). Three programs (LEA, OutFLANK, pcadapt) represented 60 percent of all submissions. The small bias toward the use of those programs could be explained by the ease to install them as R packages. The balanced software usage distribution indicated that the users were able to run the 6 programs presented during the tutorial sessions.

**Fig 2:**
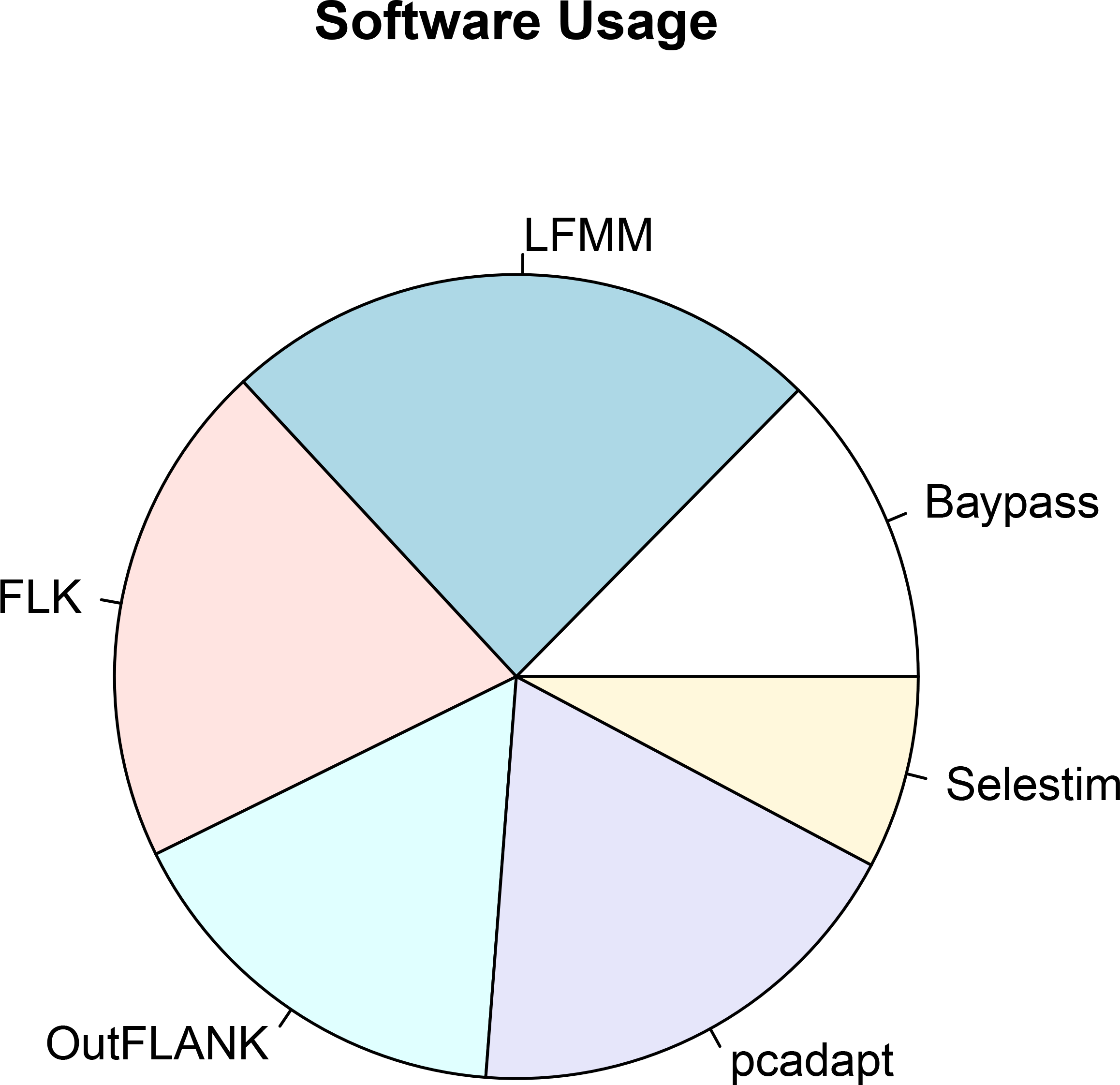
Distribution of software usage for the second and third challenges.

## Variability of scores

Candidate lists that used EA methods obtained higher scores than lists that used PD methods (Figure 3). For both PD and EA approaches and for challenges 2 and 3, the scores were highly variable (standard deviations were in the range 0.18–0.22). We also found high variability of scores when considering the distribution of scores obtained with each program separately. For the second challenge, standard deviations ranged between 0.01 and 0.34 for the six methods. For 3 programs, the scores had standard deviations around 0.20 (+/− 0.02). For the third challenge, standard deviations were between 0 and 0.22 depending on the software used, and 3 programs had a standard deviation in score around 0.20 (+/− 0.02).

Next we analyzed the scores obtained by each category of users: advanced and less-advanced users. In challenge 2 (intermediate level difficulty), the average score of advanced users was significantly greater than less-advanced users (advanced mean 0.6, less-advanced mean 0.39, t-test P = 0.05). In challenge 3 (high level of difficulty), the average score of advanced users was not significantly different than less advanced users (advanced mean 0.35, less-advanced mean 0.30, t-test P = 0.45). Similar results were obtained when we considered PD and EA methods separately, but none indicated significant differences between the two groups of users in challenge 3.

**Fig 3:**
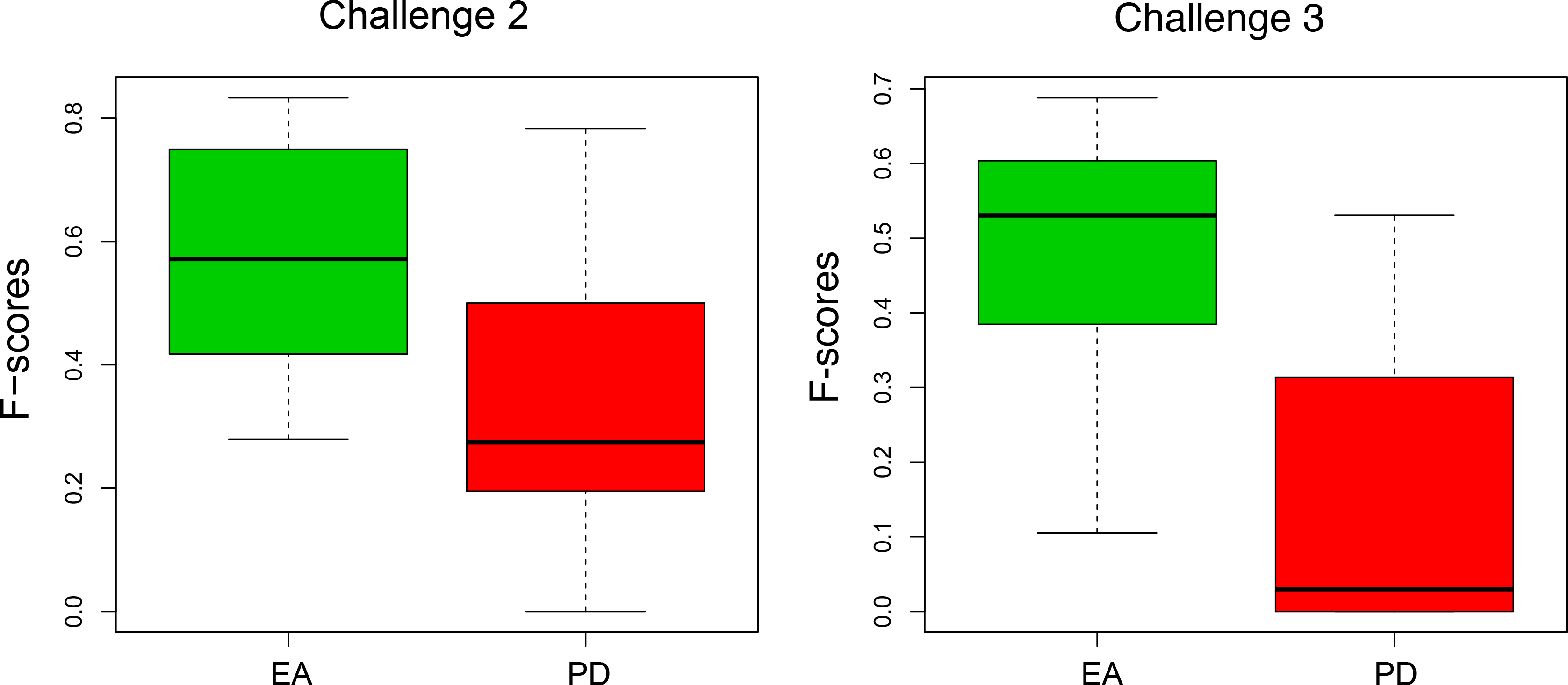
Distribution of F-scores for EA and PD methods. The left and right
panel display the results obtained for the second and third dataset, respectively. EA stands for ecological association and PD stands for population differentiation.

## Discussion

The experiment led during the summer school SSMPG 2015 showed that participants were able to learn how to use complex software for genome scans after short introductory tutorials. In challenge 1 and 3, less advanced users obtained scores comparable to those of advanced users confirming that a rapid appropriation of software based on complex statistical methods was possible. To encourage submissions, organizers delivered (symbolic) prizes to the two top-ranked user teams for the second and third challenge. Users who won the challenge prizes did not focus on a particular method, but used combinations of several methods, which is a promising direction to increase statistical power of existing methods.

## Why are the scores variable?

In this discussion, we introduce a simple probabilistic model for interpreting the variability of *F*-scores. For each adaptive locus *s*, we assume that a user discovers locus *s*, i.e., includes locus s in her/his list of candidate loci, with probability *p_s_*. We call these quantities *the detection probabilities,* and we assume that the detection probabilities depend on the level of self-declared expertise of each user. In all challenges, we observed that the users generally evaluated the number of loci under selection rather accurately. Under this condition, the expected value and the variance of the *F*-scores can be investigated theoretically (results in BOX 1)

From the theoretical results, we obtain that the variability of scores is directly related to the probabilities of correctly identifying each loci as truly adaptive. We find that the variability of F-scores is low when the challenges are either difficult or easy *(p_s_* are close to 0 or 1), and it is maximized for intermediate values of *p_s_*. For example, let us consider the results of challenge 2 for which the expected score was around .40 and the standard deviation around 0.17 (standard deviations averaged over the different software). We computed the proportion of times each adaptive locus was contained in the submitted lists, and found that the 12 truly adaptive loci spanned a range in their frequency of detection from ~0.1 to 1.0 (Figure 4). A more complex version of the probabilistic model where probabilities of detection can be different for advanced and non-advanced users (see BOX 1) reproduces the distribution of F-score for challenge 2. Assuming that probabilities of detection are reduced by 40% for nonadvanced users, that one fourth of the adaptive loci are easily detected by advanced users (*p_s_*=0.9), one half have intermediate probabilities (*p_s_*=0.65), and one fourth are difficult to detect (*p_s_*=0.3), the probabilistic model generates a distribution of *F*-scores comparable to the one obtained for challenge 2 (average F-score of 0.41 and standard deviation of 0.16).

The mathematical results in BOX 1 suggest that the variability of test performances is an inherent characteristic of the data. So, how do we interpret this variability? First, the mathematical results tell us that efforts asked to practitioners in order to reduce the variability of their scores might decrease the overall performance of tests. For example, one might recommend filtering out loci estimated to have small or medium effects on selective traits, and retain only those with strong evidence of selective signals. Such conservative practices would homogenize differences between advanced and less-advanced users and indeed decrease the variability of scores. However this strategy would also result in an increased number of false negative tests, and reduce the overall value of the expected score.

The previous example illustrates that the expected *F*-score could be increased if users increase their expertise (and therefore the probability to detect loci of medium effects). Although increasing user expertise is a desirable point, the equations tell us that the variability of scores would not necessarily be decreased. For example, assume there are 12 SNPs of medium effect under selection and the probability of detection of each locus is of 50% for advanced users and of 5% for less advanced users. A group with 5% advanced users and 95% less-advanced users would have an expected F-score of 0.07 and a standard deviation in F-scores of 0.12. Now, assume that all users become advanced; although the expected F-score of the advanced group is higher (0.5), the standard deviation in F-scores is also increased, as it is equal to 0.14. This example shows that reducing the variance of F-scores is not a desirable objective, because in some cases increasing user expertise can result in an increase in the variance of *F*-scores.

The uneven difficulty of detecting adaptive loci, which explains the variability of scores, arises from methods, but also from the decisions made by users regarding model parameters, test calibration and the choice of a false discovery rate in statistical tests. To illustrate how much user decisions can influence the variability of scores, we re-ran two programs, pcadapt and LFMM, on challenges 2 and 3 (using K = 3 in both programs). The runs of each program resulted in well-calibrated *P*-values for each data set (FranÇois et al. 2016). Then we built lists of candidate loci by using an FDR control algorithm (Benjamini & Hochberg 1995). The algorithm requires that an expected level of FDR is specified by the user, and uses the expected level of FDR to determine a list of candidates. To simulate users’ decisions, we sampled expected levels of FDR according to a beta distribution of mean 0.05 and standard deviation 0.047. The distribution of scores from the created lists had standard deviations of 0.14 and 0.09 for dataset 2 and dataset 3 respectively. These results provide evidence that decisions about how to use the program outputs can generate large variability of scores as observed in Figure 3.

**Fig 4:**
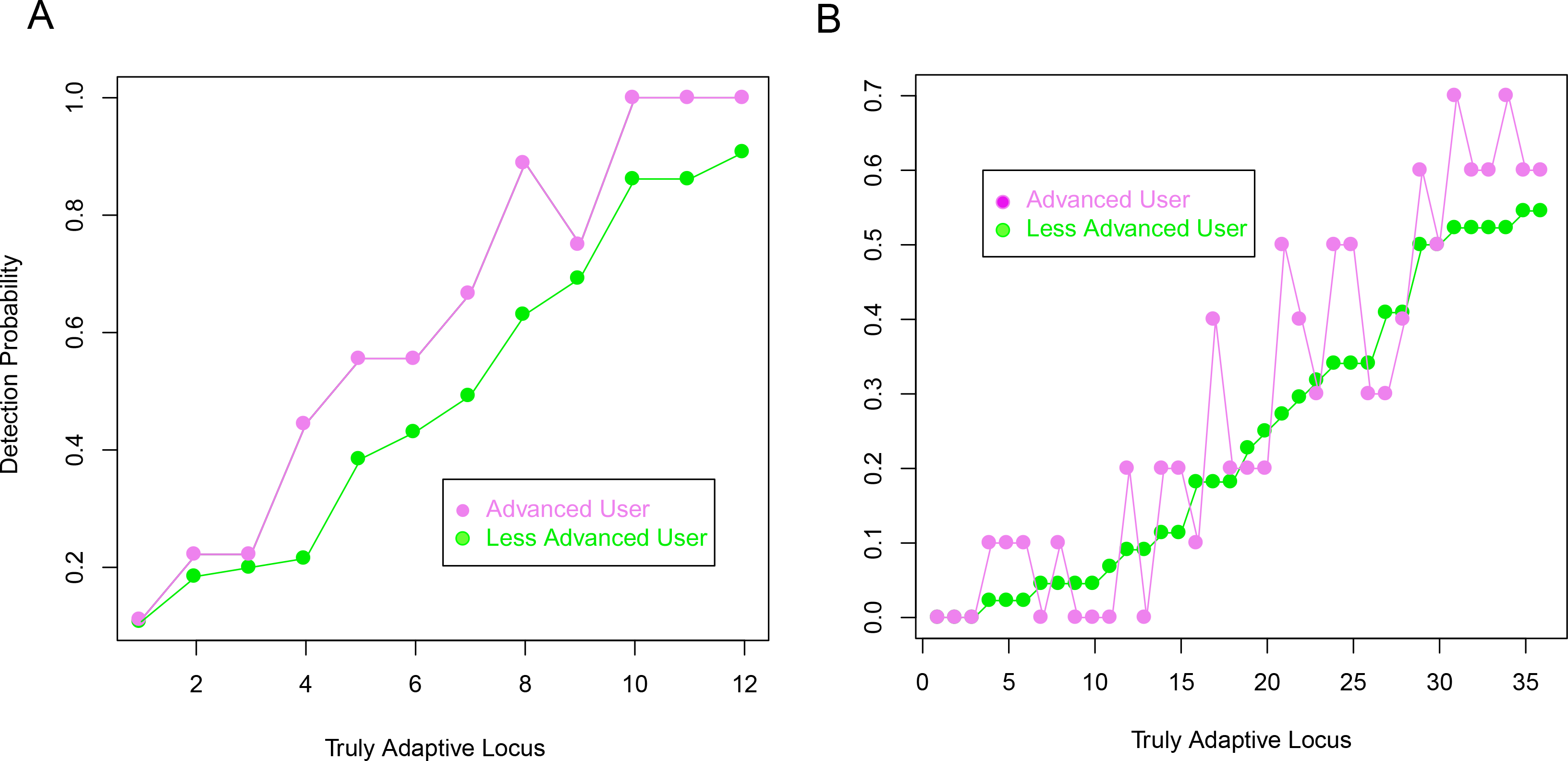
Probabilities of detection of adaptive loci. For each locus, we counted the proportion of times it was contained in a submitted list. A) Second challenge: probabilities of detection were larger for advanced users than for less-advanced users. B) Third challenge: No obvious differences in detection probabilities when comparing advanced and non-advanced users.

## Conclusions

The obvious lesson of the SSMPG15 experiment was to promote the usage of powerful statistical methods and simultaneously improve the expertise of their users. The first action is the goal of current methodological developments of genome scans for selection, which should always be accompanied by clear and practical user guides. The second action requires that practitioners develop their own statistical and computer skills to follow the rapid development of complex methods. To provide a training opportunity, the website containing the data presented during SSMPG 2015 as well as additional new datasets is now publicly available (https://ssmpg-challenge.imag.fr/). On the website, users can submit lists of candidate loci that will be scored and compared to the scores obtained by previous users of the website.

## Box 1: Explaining the variability of *F*-scores

To explain our results and final recommendations, we introduce a simple probabilistic model for interpreting the variability of *F*-scores. For a given challenge, we assume that the unknown list of truly adaptive loci contains *m* elements. The values of *m* were equal to *m* = 12 in the first and second challenge and *m* = 36 in the third challenge. For each adaptive locus, *s*, we denote by *p_s_* the probability (power) that a user discovers locus *s*, i.e.,includes this locus in her/his own list of candidate loci. At SSMPG15, the submitted user lists were approximately of length *m*, meaning that the users correctly evaluated the number of loci under selection in the simulated datasets. Under this hypothesis, we can compute the expected value of the *F*-score, E[*F*]. We obtain that E[*F*] is equal to (p_1_+…+p_m_)/m. Assuming that the tests are independent, the variance of *F*–scores, Var(*F*), is equal to (p_1_(1–p_1_)+…+p_m_(1–p_m_))/m^2^. From these results, we obtain that the variability of *F*-scores is directly related to the probabilities to correctly identify each loci as truly adaptive. The variability of F–scores is low when the challenges are difficult or easy (*p_s_* close to 0 or 1), and it is maximized for intermediate values of *p_s_*.

Accounting for the self-reported expertise of each user, we defined two categories of users: advanced ones, A, and less-advanced ones, *A'.* In this context, the statements about the expected value and the variance of *F*-scores can be refined as follows. Consider π_A_ the proportion of advanced users, and write 1 - π_A_ for less-advanced users. For a truly adaptive locus s, there is a conditional probability, *p_sA_*, for locus *s* to be correctly identified by an advanced user, and there is a lower probability, *p_sA,_,* for *s* to be correctly identified by a less-advanced user. The expected value of F-scores can be computed as follows 
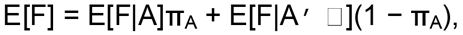
where the conditional expectation E[F|A] is equal to (p_1A_+…+p_mA_/m, and where the formula for E[F|A'] is similar. In other words, the conditional expectations are computed by averaging the conditional probabilities over all adaptive loci. The variance of F-scores decomposes as follows
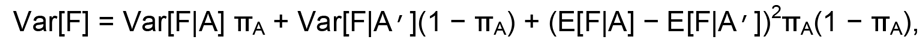
where Var[F|A] is equal to (p_1A_(1– p_1A_)+…+p_mA_(1– p_mA_))/m^2^, and the formula for Var[F|A'] is similar. In the discussion, we use the formulae for the mean E[F] and the variance Var[F] to measure the effect of filtering strategies and of increasing user expertise on the distribution of *F*-scores.

## Acknowledgments

We acknowledge Kevin Caye, Thomas Dias-Alves, and Keurcien Luu for developing the website, and Frédéric Guillaume for providing support with the simulations in NEMO. We would like to thank our colleagues who were instructors during the SSMPG summer school (Mathieu Gautier,Bertrand Servin, and Renaud Vitalis) as well as the attendees of the SSMPG 2015 summer school.

## Supplementary Figure 1

Schematic temporal evolution of the species range used to simulate data.Populations 1-3 and populations 15-17 correspond to the two initial glacial refugees. Populations 6,9, and 10 correspond to the summit that was possible to colonize after generation 3,000. The diameters of the circles increase with increasing effective population size that range from N=500 to N=2500. Colors on the landscape represent the phenotypic optimum for the trait, ranging from an optimum of −3 (dark green) to 3 (white).

## Supplementary Figure 2

Parameter settings used in the simulations of the three different challenges.

